# FlashPCA: fast sparse canonical correlation analysis of genomic data

**DOI:** 10.1101/047217

**Authors:** Gad Abraham, Michael Inouye

**Affiliations:** Centre for Systems Genomics, School of BioSciences, University of Melbourne, Parkville 3010, VIC, Australia.; Department of Pathology, Faculty of Medicine, Dentistry, and Health Sciences, University of Melbourne, Parkville 3010, VIC, Australia.

## Abstract

**Summary:** Sparse canonical correlation analysis (SCCA) is a useful approach for correlating one set of measurements, such as single nucleotide polymorphisms (SNPs), with another set of measurements, such as gene expression levels. We present a fast implementation of SCCA, enabling rapid analysis of hundreds of thousands of SNPs together with thousands of phenotypes. Our approach is implemented both as an R package flashpcaR and within the standalone commandline tool flashpca.

**Availability and implementation:** https://github.com/gabraham/flashpca

## 1 Introduction

Canonical correlation analysis (CCA) is a well-known statistical approach for multivariate analysis of two datasets (Hotelling, 1936). In the context of large-scale genomic and multi-omic analyses, CCA can prove useful in identifying relationships amongst complex data, for example single nucleotide polymorphisms (SNPs) and gene expression levels. Approaches that consider one SNP at a time together with multiple phenotypes have been shown to increase power to detect QTLs over the simpler but commonly utilised single-SNP/single-phenotype approach (Ferreira et al., 2009; Inouye et al., 2012), particularly when analysing correlated phenotypes that are modulated by the same genetic variants.

Analysis of multiple SNPs simultaneously is an attractive extension of the single-SNP multiple-phenotype approach, however, standard CCA is not well-defined when the number of samples is lower than the number of SNPs or phenotypes (*n*< *min*{*p*,*m*}). One solution is Sparse CCA (SCCA) (Witten et al., 2009a,b; Parkhomenko et al., 2009), an *L*_1_-penalised variant of CCA which allows for tuning the number of variables that effectively contribute to the canonical correlation, thus making the problem well-defined. Owing to the induced sparsity, SCCA can be useful for identifying a small subset of SNPs and a small subset of the phenotypes exhibiting strong correlations. However, the rapidly increasing size and coverage of genotyping arrays (exacerbated by genotype imputation), together with the availability of large phenotypic datasets (transcriptomic, metabolomic, and others; e.g., Bartel et al. 2015); The GTEx Consortium (2015)), makes it challenging to perform such analyses using existing tools.

We have developed an efficient implementation of SCCA that is capable of analysing genome-wide SNP datasets (1 million SNPs or more) together with thousands of phenotypes, as part of the tool flashpca (Abraham et al., 2014). The tool is implemented in C++ using the Eigen 3 numerical library (Guennebaud et al., 2010), as well as an R interface (package flashpcaR) based on RcppEigen (Bates et al., 2013).

Here, we compare the SCCA implementation in flashpcaR and flashpca with a widely-used implementation (PMA, by Witten et al. (2013)), and demonstrate the substantial improvements in speed of our tool, allowing for large analyses to be performed rapidly.

## 2 Methods

In standard CCA, we assume that we have two matrices **X** (*n* × *p)* and **Y** (*n* × *m*), measured for the same *n* samples. We further assume that both **X** and **Y** have been column-wise standardised (zero mean, unit variance). For a single pair of canonical variables *a* and *b*, CCA involves solving the problem

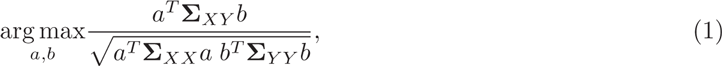

where *Σ*_*XX*_ and *Σ*_*YY*_ are the covariance matrices of **X** and **Y**, respectively, and *Σ*_*XY*_ is the covariance matrix of **X** and **Y**. The solution is given by the singular value decomposition (SVD) of 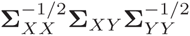, with 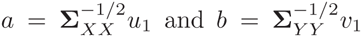, where *u*_1_ and *v*_1_ are the first left and right singular vectors, respectively.

SCCA is typically used for high-dimensional data, where a useful assumption is that the columns of **X** and **Y** are uncorrelated, i.e., *Σ*_*XX*_ = *Σ*_*YY*_ = **I** (Parkhomenko et al., 2009), hence, *a = u* and *b = v.* Thus, SCCA involves solving another form of CCA,

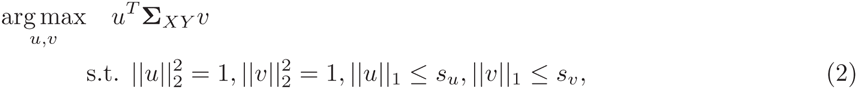

where *u* and *v* are the left and right canonical vectors, respectively, and *s*_*u*_ and *s*_*v*_ are constraints on the *L*_1_ norms of these canonical vectors.

The problem can be converted into the penalised (Lagrangian) form and solved using iterative soft-thresholding (Parkhomenko et al., 2009). Unlike standard CCA, SCCA is well-defined even when *n<m*in{*p*,*m*}, and induces sparse canonical vectors, depending on the choice of *L*_1_ penalties (higher penalties lead to higher sparsity). The optimal set of penalties can be found via cross-validation: the data (both **X** and **Y**) are split into training and test sets, SCCA is run on the training set (**X**_train_, **Y**_train_) using a 2D grid of penalties, and the pair of penalties that produce the highest correlations in the test set, Cor(**X**_test_*u*, **Y**_test_*v*), are selected.

## 3 Results

We utilised the HapMap3 phase III genotypes (International HapMap 3 Consortium, 2010), together with gene expression data of709 individuals (Stranger et al., 2012). After quality control (see Supplementary Material) and taking the intersection of SNPs across the populations, the data consisted of 709 individuals, 973,983 SNPs, and 18,379 gene expression probes.

We first confirmed that `flashpcaR::scca` produced models comparable with `PMA::CCA`, by comparing the results in cross-validation on HapMap3 genotypes together with simulated gene expression levels (Supplementary Material). Next, to further assesss the relative speed improvement using real-world data, we used subsets of the HapMap3 genotypes and real gene expression levels (Stranger et al., 2012) and compared the runtime of: (i) `PMA::CCA` (R package), (ii) `flashpcaR::scca` (R package), and (iii) flashpca (commandline tool). Whereas both `PMA::CCA` and `flashpcaR::flashpca` are bound by the memory limitations of R, the commandline tool flashpca allows much larger analyses; hence we also ran larger analyses: chromosomes 1-2, 1-3, …, 1-22, up to all 973,983 SNPs. Figure 1 shows that `flashpcaR::scca` was 3-12 × faster than `PMA::CCA`, with an analysis of 75,000 SNPs and 10,000 gene expression levels completing in 5s and 50s, respectively. The commandline flashpca was faster than `PMA::CCA` as well, and completed an analysis of 709 individuals, 973,983 SNPs and 18,379 gene expression levels in median wall time of ~47s (including all overheads), using ~10GiB of RAM. Note that runtime for the commandline `flashpca` includes all steps such as loading data into RAM, unlike the R version where the data is pre-loaded into R. Performing cross-validation over a grid of penalties will increase these times, and we recommend parallelisation over several cores (Supplementary Material).

**Figure 1:**
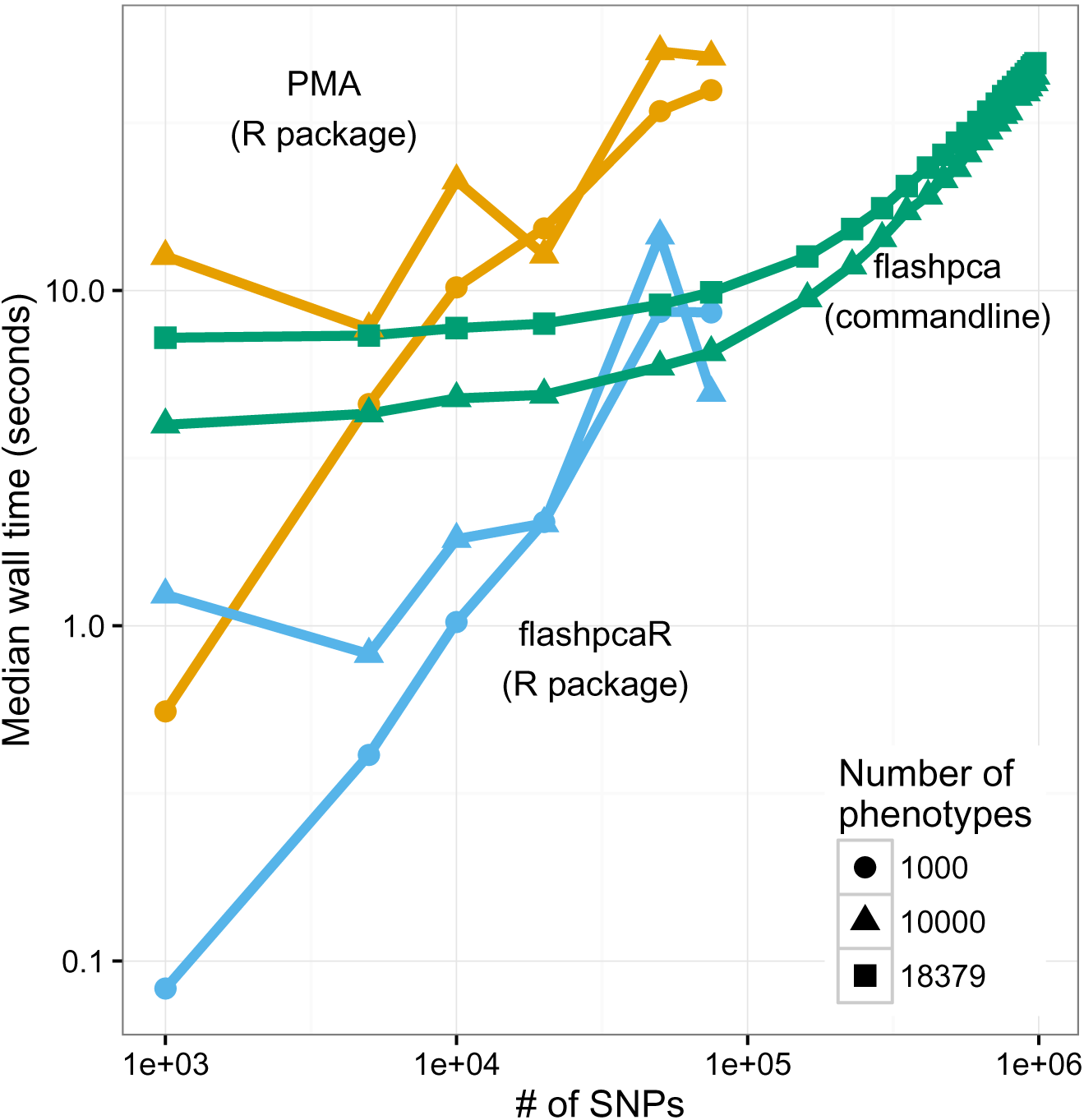
Timing (median of 30 runs) of SCCA implemented in (i) the `flashpcaR` (`R` package) and (ii) flashpca (stand-alone commandline tool), compared with `PMA`, using subsets of the HapMap3 dataset with gene expression levels as phenotypes. The stand-alone flashpca timing includes data loading into memory.

## 4 Conclusion

`flashpca` provides a fast implementation of sparse canonial correlation analysis, making it possible to rapidly analyse high dimensional datasets. SCCA is available in the R package `flashpcaR`, which enables analysis of metabolomic, transcriptomic, or any other quantitative set of measurements. The commandline version is targeted at SNP/phenotype data, enabling large QTL analyses of >1 million SNPs and thousands of phenotypes, that would otherwise be too large to fit within R.

